# Visualizing multi-protein patterns at the synapse of neuronal tissue with DNA-assisted single-molecule localization microscopy

**DOI:** 10.1101/2021.02.23.432306

**Authors:** Kaarjel K. Narayanasamy, Aleksandar Stojic, Yunqing Li, Steffen Sass, Marina Hesse, Nina S. Deussner-Helfmann, Marina S. Dietz, Maja Klevanski, Thomas Kuner, Mike Heilemann

**Affiliations:** Department of Functional Neuroanatomy, Institute for Anatomy and Cell Biology, Heidelberg University, Germany; Institute of Physical and Theoretical Chemistry, Goethe University, Frankfurt am Main, Germany

**Keywords:** single-molecule localization microscopy, super-resolution microscopy, DNA-PAINT, neuronal synapse, multiplexing

## Abstract

The development of super-resolution microscopy (SRM) has widened our understanding of biomolecular structure and function in biological materials. Imaging multiple targets within a single area would elucidate their spatial localization relative to the cell matrix and neighboring biomolecules, revealing multi-protein macromolecular structures and their functional co-dependencies. SRM methods are, however, limited to the number of suitable fluorophores that can be imaged during a single acquisition as well as the loss of antigens during antibody washing and restaining for organic dye multiplexing. We report the visualization of multiple protein targets within the pre- and postsynapse in 350-400 nm thick neuronal tissue sections using DNA-assisted single-molecule localization microscopy. Using antibodies labeled with short DNA oligonucleotides, multiple targets are visualized successively by sequential exchange of fluorophore-labeled complementary oligonucleotides present in the imaging buffer. The structural integrity of the tissue is maintained owing to only a single labelling step during sample preparation. Multiple targets are imaged using a single laser wavelength, minimizing chromatic aberration. This method proved robust for multi-target imaging in semi-thin tissue sections, paving the way towards structural cell biology with single-molecule super-resolution microscopy.

## Introduction

Super-resolution microscopy has revolutionized our understanding of cell biology. Single-molecule localization microscopy (SMLM) is one branch of super-resolution microscopy, which employs photoswitchable or transiently binding fluorophore labels and has demonstrated a near-molecular spatial resolution (Sauer and Heilemann 2017) allowing molecular quantification (Dietz and Heilemann 2019). A further exciting development was the integration of short DNA oligonucleotides into the concept of SMLM, as realized in DNA point accumulation in nanoscale topography (DNA-PAINT) (Jungmann et al. 2010). The short oligonucleotides act as transiently hybridizing pairs, with one coupled to a target protein (the “docking strand”, attached to e.g. an antibody) and a second carrying a fluorophore (the “imager strand”) suspended in the imaging buffer. The transient hybridization of both oligonucleotides generates a temporally short and spatially localized signal, which at low concentration of imager strands is recorded as a single-molecule emission event. A particular strength of DNA-PAINT is that multi-color imaging is not limited by the number of fluorophores that can be separated by their emission spectra, but instead the “color” is encoded into the DNA sequence of the pair of docking and imager strand utilized in consecutive imaging rounds. Implementing an experimental protocol that exchanges imager strands in the buffer solution allows for imaging of more targets than if discrimination occurs on the basis of emission spectra (Jungmann et al. 2014). Multiplexing and the excellent spatial resolution achieved with DNA-PAINT is now beginning to evolve as a tool in cell biology (Strauss and Jungmann 2020; Schröder et al. 2020; Harwardt et al. 2020).

The next important step in the application of super-resolution microscopy to cell biology is to visualize the nano-architecture of proteins in the functional context, which demands for super-resolution imaging in tissue and multiplexed imaging of many proteins in the same sample. SMLM imaging of 15 protein targets in cells and tissue was recently achieved using multiple rounds of antibody labeling and fluorophore staining (Klevanski et al. 2020). Here, we demonstrate the integration of DNA-PAINT for super-resolution imaging of structurally preserved neuronal brain tissue from rats, and we achieve a spatial resolution of better than 30 nm. We demonstrate multiplexed imaging of four targets using only one excitation laser light source, which eliminates chromatic aberration. Furthermore, we integrate recent developments in DNA-PAINT labels that allow for faster imaging (Strauss and Jungmann 2020). In short, we established an experimental pipeline for robust and fast super-resolution imaging of proteins in structurally preserved tissue that achieves near-molecular spatial resolution and enables the ultrastructural investigation of protein assemblies in their native environment.

## Results

We employed Exchange PAINT (Jungmann et al. 2014) for super-resolution imaging of multiple protein targets in neuronal tissue. Using this technique, four proteins were immunolabeled simultaneously, thereby maintaining low sample preparation time while obtaining an information-rich dataset. In a first experiment, α-tubulin, mitochondria (TOM20), microtubule-associated protein 2 (MAP2), and vesicular glutamate transporter (VGLUT1) were labelled with primary antibodies (Ab) and secondary Ab conjugated to DNA docking strands (P1, P5, R1, or R4; see Methods) (Figure 1).

**Figure 1:**
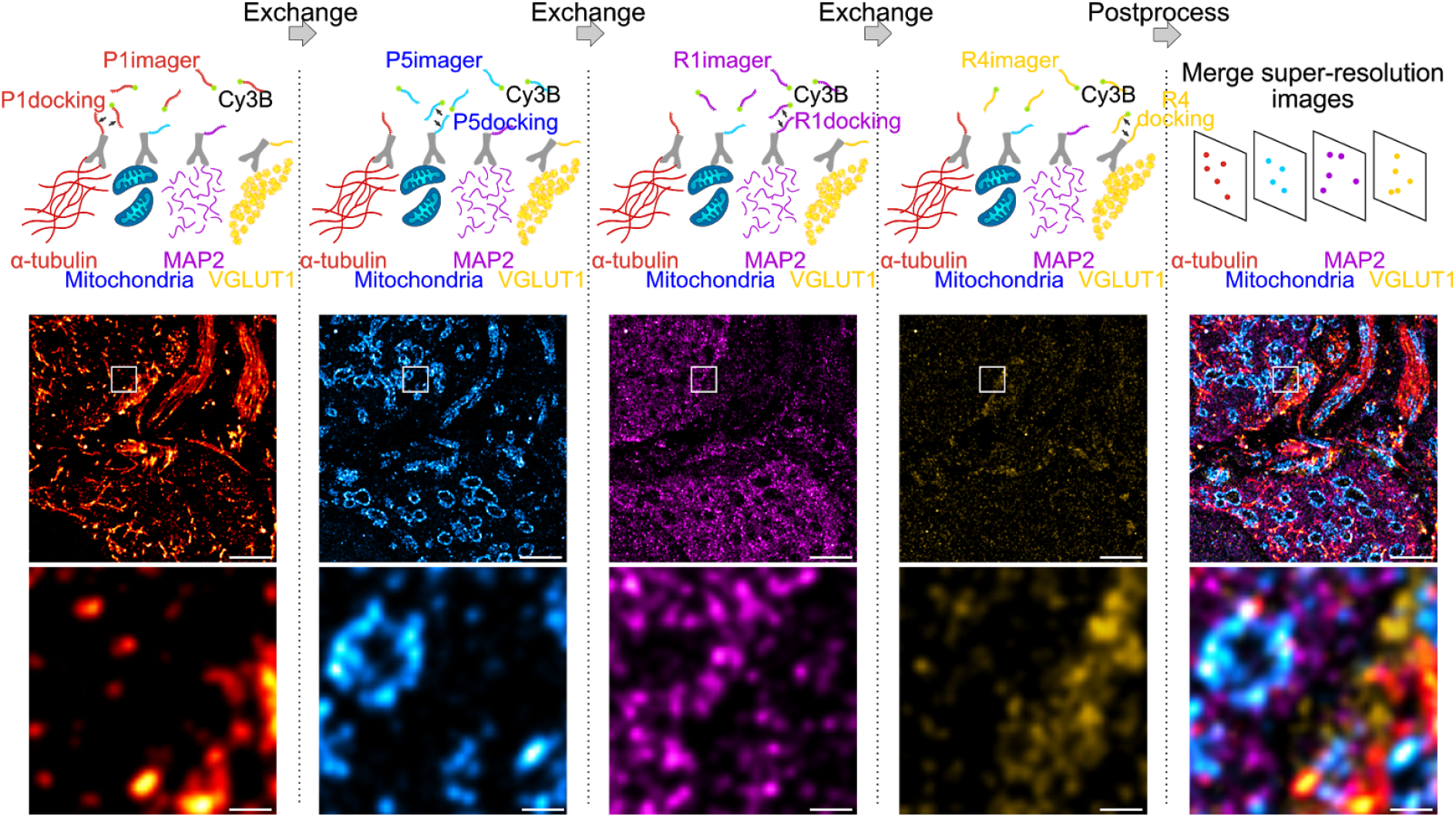
Exchange DNA-PAINT of four targets imaged sequentially. Four protein targets in tissue were labelled with primary antibodies and their corresponding secondary antibody-docking strand conjugate (P1, P5, R1, or R4). The Cy3B labelled imager strands were imaged sequentially by strand type with wash steps between each imaging round. All SMLM rendered images depicting each target were merged to obtain a multi-protein super-resolved image. Scale bar 1 µm (top) and 0.1 µm (bottom).

The protocol for sequential DNA-PAINT imaging started by adding P1 imager strands into the buffer and imaging α-tubulin in the first round, followed by washing away the strands and replacing them with P5 imager strands for mitochondrial imaging. This set of steps was repeated with R1 and R4 strands until all labelled proteins were imaged within the same region of interest (ROI). Each set of frames was rendered individually and merged together using fiducial markers to obtain an overlay of four protein targets organized within tissue (see Methods).

This method was implemented to study the structure and organization of proteins in semi-thin neuronal tissue sections, specifically within the medial nucleus of the trapezoid body (MNTB) region which contains specialized cells known as the calyx of Held (Figure 2a, inset). These calyces are giant presynaptic terminals (grey) partially enveloping the postsynaptic principal cell (purple) with finger-like protrusions. Each calyx contains hundreds of active zones (AZs) for glutamatergic synaptic transmission (Dondzillo et al. 2010; Sätzler et al. 2002). A transverse section of the calyx of Held reveals the soma of the principal cell and presynaptic endings distributed around the edges, exposing the AZs of the synaptic contact. α-tubulin, mitochondria, MAP2, and VGLUT1 were stained with the Ab-DNA conjugate and imaged with Exchange PAINT (Figure 2a). The image shows several principal cells enveloped by the presynaptic calyx of Held, two of them fully visible within the tissue matrix (stippled lines), with one sectioned across the nucleus, as well as axons and capillaries (dotted line). MAP2 is commonly used as a neuronal marker as it selectively labels neuronal cells, specifically the cytoplasm of the soma and dendrites (Sarnat 2013). VGLUT1 is a marker for synaptic vesicles (SVs), which are concentrated in the presynaptic terminal of the calyx. Regions with interesting morphological and organizational protein distribution are magnified in figures 2 i – iv, representing the co-organization between tubulin (red) and mitochondria (cyan) within morphologically distinct structures. Figure 2 i & ii are the transverse and cross-sections of axons, respectively, which show the parallel organization of tubulin filaments along the length of the axon or the circular arrangement of tubulin within an axon bundle. Mitochondria within the axons are thin, elongated structures sandwiched between tubulin filaments and are distributed randomly along and across the axon bundle. The protein organization seen here is in line with the knowledge that tubulin filaments (microtubules) play a role in mitochondrial transport along axons to the presynaptic terminals where they are needed to maintain continuous synaptic transmission (Zorgniotti et al. 2021; Verstreken et al. 2005).

**Figure 2:**
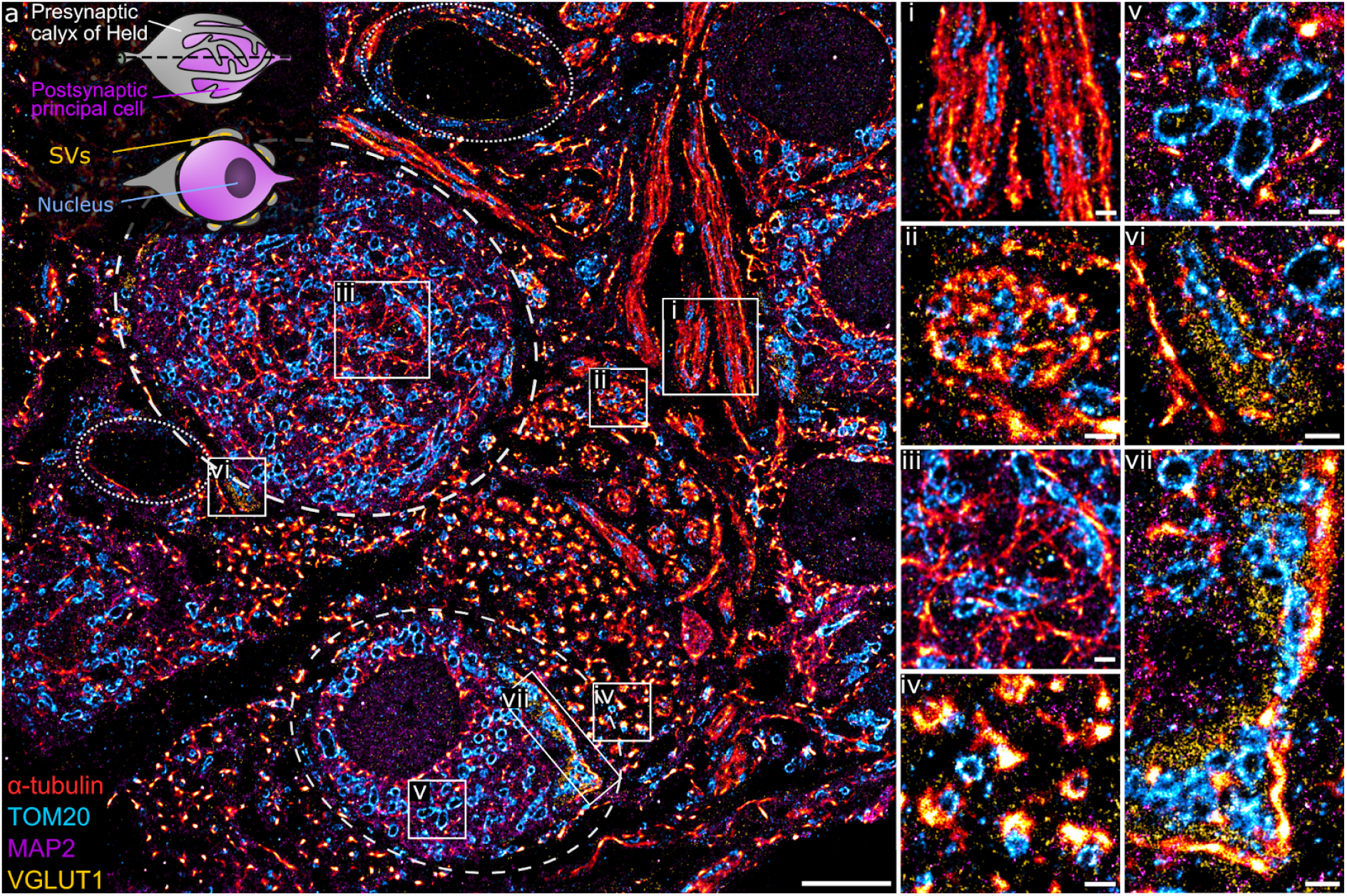
(a) A 4-target overlay DNA-PAINT image of MNTB tissue with two MNTB synapses (stippled lines), capillaries (dotted lines), and a graphical representation of the calyx of Held (inset). (i -vii) Magnification of regions within the primary image (a) showing different protein morphologies and organization of tubulin, mitochondria, MAP2, and VGLUT1 within the MNTB. Scale bar 5 µm (a) and 0.5 µm (i – vii).

Apart from axons, tubulin and mitochondria are also co-organized in other parts of the neural network. Figure 2iii shows the organization between tubulin and mitochondria within the soma of the principal cell. Here, tubulin filaments appear as short, thin fibrils without a distinct organizational pattern. Similarly, mitochondria show random arrangement within the soma. MAP2 clearly labels the soma of principal cells with larger and oval shaped mitochondria embedded within the matrix (Figure 2v). Another morphologically distinct structure of tubulin is observed next to the smaller MNTB synapse. Here, tubulin forms dense, small bundles and each bundle is organized tightly with 1-2 mitochondria (Figure 2iv).

Figure 2 vi & vii show presynaptic compartments of the calyces containing SV clusters (yellow) next to the principal cell. A feature of interest is the proximity of SVs to tubulin, which can be found as punctate structures embedded on the synaptic site (Fig. 2vi) or bordering the outer edge of the SV cluster (Fig. 2vii). The close proximity of tubulin and SVs has been documented before (Piriya Ananda Babu et al. 2020) and function in the transport and regulation of synaptic vesicle precursors to the presynaptic terminal. Furthermore, mitochondria localized in between SVs in the presynapse are morphologically more compact and dense compared to those in the principal cell.

Next, we characterized the image quality using experimental parameters used for SMLM data (Sauer and Heilemann 2017). We determined the localization precision and the spatial resolution achieved with the different imager strands used in the Exchange PAINT experiment, i.e. P1, P5, R1, and R4. The P1 and P5 strands were among the first DNA sequences used in DNA-PAINT and hybridized into a duplex of 9 nucleotide base pairs (Schnitzbauer et al. 2017). The R1 and R4 strands contained repeated and concatenated sequences that allowed the hybridization of multiple imager strands onto one docking strand increasing the frequency of events (Strauss and Jungmann 2020). The localization precision of events was calculated from the nearest neighbor value (Endesfelder et al. 2014) (Figure 3a) and the lowest localization precision value obtained was 3 nm with P5 strands. Median values recorded for all four strands were below 5 nm. The spatial resolution obtained for the four imager strands was determined by a decorrelation analysis (Descloux, Grußmayer, and Radenovic 2019) which reported median values around 25 nm, and the highest resolution achieved was 21 nm for the P5 strand (Figure 3b). Interestingly, the lowest localization precision recorded here did not correspond to the highest image resolution.

**Figure 3:**
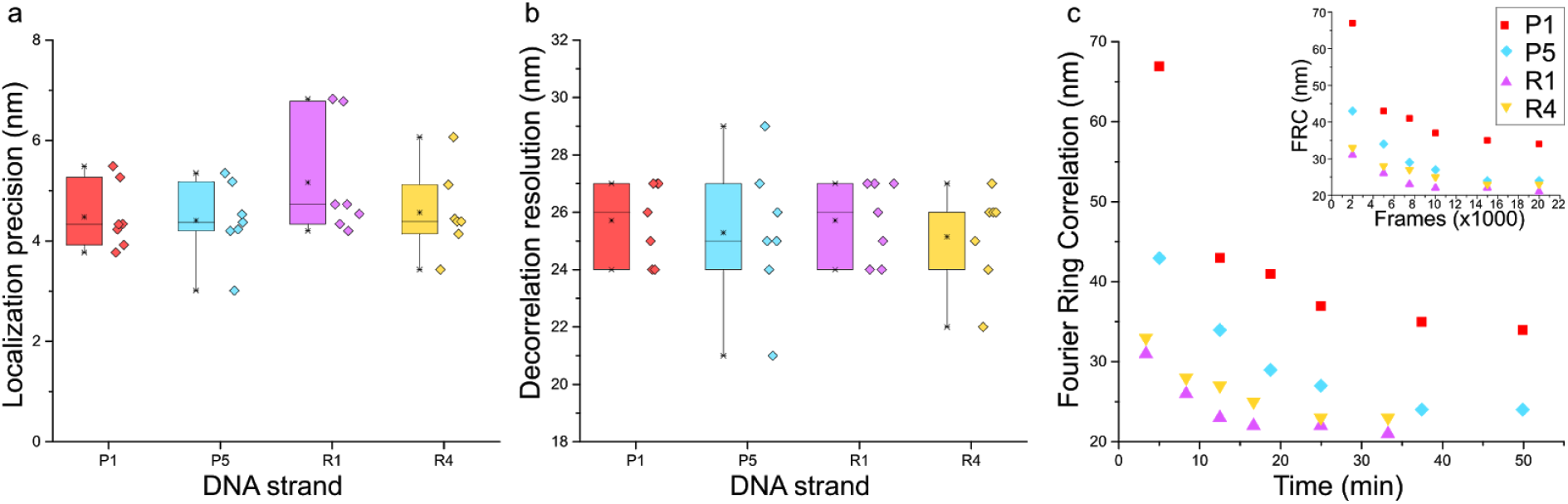
Comparison between P1, P5, R1, and R4 DNA-PAINT strands for (a) localization precision by nearest neighbour analysis (Endesfelder et al. 2014) and (b) rendered image resolution by decorrelation analysis; n = 7. (c) Fourier Ring Correlation (FRC) resolution trend of the four strands over image acquisition time and FRC over number of frames (inset); n = 1.

Although there was no apparent difference in the localization precision and resolution between the P strands and R strands, a marked advantage of the R strands was the shorter acquisition time required during imaging and increased frequency of binding between imager and docking strands, which was reported to reduce the imaging time (Strauss and Jungmann 2020). We sought to quantify this using Fourier Ring Correlation (FRC) analysis (Nieuwenhuizen et al. 2013) by calculating the resolution of images formed over time. Each super-resolved image was reconstructed from 20 000 frames with an integration time of 150 ms (P1 and P5) or 100 ms (R1 and R4), respectively. Figure 3c shows that the FRC curve plateaued before imaging time was complete, therefore all images were able to achieve maximum resolution at 20 000 frames. Saturation of resolution was calculated at 95% of the lowest resolution value achieved for each image. Indeed, both R strands were able to achieve maximum resolution faster than P strands, with R1 and R4 at 17 and 20 mins, and P1 and P5 at 37 and 34 mins, respectively. The reduction in imaging time by 15 – 20 mins, and comparable localization precision and resolution make the R strands suitable for faster Exchange PAINT imaging of multiple targets.

We next sought to apply Exchange PAINT to visualize a key component of the synaptic architecture – the AZ. Here, synaptic scaffold proteins Bassoon and Homer that delineate the active zone and postsynaptic density (PSD) were imaged in MNTB tissue to observe their distribution. The presynaptic region was identified using the VGLUT1 marker for SVs and the postsynaptic area using the neuronal marker MAP2. Multiple Bassoon (AZ) and Homer (PSD) structures represent synaptic contacts formed by the calyx and principal cell (Figure 4a i-iii). Bassoon is located on the inner presynaptic border, defined here by the inner edge of the VGLUT1 band, and Homer is juxtaposed against Bassoon and found on the edge of the MAP2 signal (Figure 4a iv-vi & b). Magnified images of Bassoon and Homer show highly resolved edges and a defined space in between, partially reflecting the presence of the synaptic cleft, as well as curved (Figure 4a iv&v) or linear morphologies (Figure 4a vi) of the AZ and PSD.

**Figure 4:**
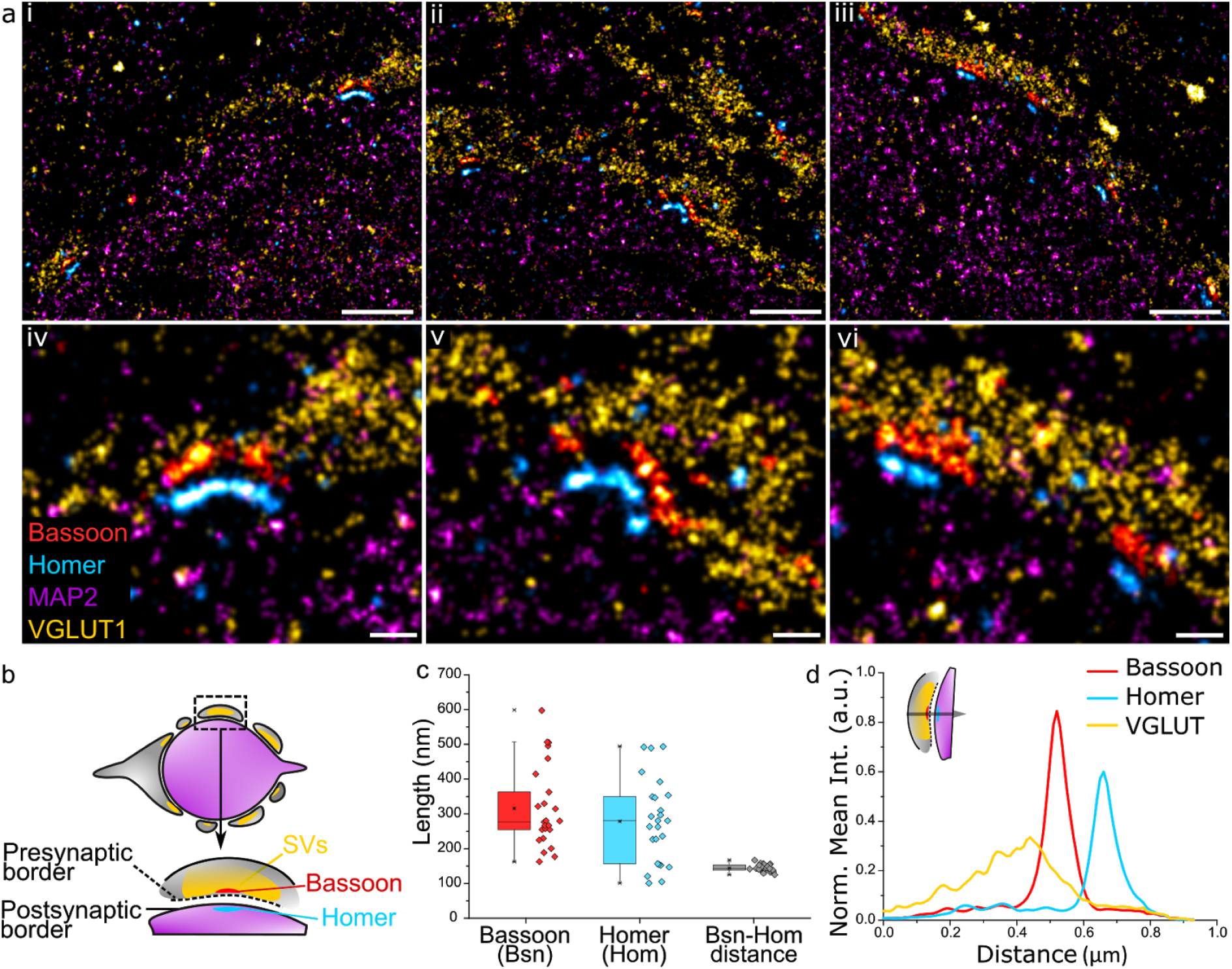
(a) Four-target images of (i -iii) the organization of multiple Bassoon and Homer structures sandwiched between VGLUT1 (SV) and MAP2 (microtubules) along the pre-and postsynaptic border of the calyx of Held. (iv -vi) Magnification of the AZ-PSD interface with aligned Bassoon and Homer structures showing linear or curved morphologies. (b) Graphical representation of a trans-section of a calyx of Held principal cell (purple) surrounded by the presynaptic cell (grey) and the organization of Bassoon, Homer, and SVs. (c) Quantification of the length of Bassoon-or Homer-positive areas, and the distance between Bassoon to Homer; n = 25. (d) Line profile of 2D spatial organization of protein density based on fluorescence intensity from VGLUT1 to Bassoon to Homer; n = 16. Scale bar 1 µm (i – iii) and 0.2 µm (iv – vi).

The profile views of Bassoon and Homer were measured lengthwise (Figure 4c) and had a comparable median length of 277 nm and 281 nm respectively (Bassoon mean = 316 nm, SD = 117; Homer mean = 278 nm, SD = 120). Of considerable interest in studies of synaptic specializations is the distance between Bassoon and Homer. These scaffold proteins are located below their respective synaptic borders, therefore, unlike the synaptic cleft which has a distance of only 28 ± 9 nm in the calyx (Sätzler et al. 2002), are spaced far enough apart to be easily resolved using super-resolution light microscopy. The Bassoon-Homer distance was found to fall into a narrow distribution range with median 143 nm (mean = 144 nm, SD = 10 nm; Figure 4c). This distance is reflected in the intensity line profiles of Bassoon and Homer (Figure 4d). This value is in good agreement with previous SMLM studies reporting values of ∼150 ± 20 nm in brain tissue (Dani et al. 2010) and ∼165 ± 9 nm in neuronal cultures (Glebov et al. 2016).

The relative localization of VGLUT and Homer to Bassoon was determined by measuring the fluorescence intensity profile of the corresponding proteins within the AZ from the presynaptic terminal towards the principal cell (Figure 4d & inset). The 2D line profile shows defined Bassoon and Homer peaks with respective widths of 82 nm and 85 nm at FWHM (Gaussian fitting). SVs are found to be anchored at higher density closer to Bassoon. SVs function in the release of neurotransmitters at the presynaptic AZ, hence are present in high density on the presynaptic membrane. While the exact function of Bassoon is still unknown, it was shown to play a role in short-term SV replenishment during neurotransmission (Dani et al. 2010; Parthier, Kuner, and Körber 2018; Hallermann et al. 2010) and SV tethering to the AZ (Mukherjee et al. 2010), thus accounting for the colocalization of SVs and Bassoon.

## Discussion

Studies in structural biology require imaging in greater spatial resolution and to observe proteins in their native environment. One of the challenges in imaging neuronal structures is studying the precise organization of proteins within a dense spatial matrix as well as their relative localization to other neuronal proteins. To this end, super-resolution microscopy has been used as a tool due to its ability to resolve structures in the nanoscale and image multiple targets to obtain an overview of protein arrangement within neurons (Colnaghi et al. 2020); (Kubo et al. 2019), and has shed light on disease pathologies within dense structures (Shahmoradian et al. 2019).

SMLM methods such as STORM and Bayesian blinking and bleaching (3B) have been used to study the organization of proteins in the AZ (Glebov et al. 2016; Dani et al. 2010). However, the number of spectrally distinct fluorophores that can be used for photoswitching and which chromatic aberration can be corrected are limited, which prevents the imaging of more than three structures at a time. To overcome this, super-resolution imaging with dSTORM (Heilemann et al. 2008) was accomplished by sequential staining realized via bleaching, elution, and restaining using antibodies or other labels against 16 protein targets to obtain an overview of protein distribution within the calyx of Held (Klevanski et al. 2020). The signal density of a target protein can be enhanced by implementing multiple rounds of labeling and imaging (Venkataramani et al. 2018). An alternative solution to visualize protein targets in super-resolution microscopy is the integration of DNA-based protein labels (e.g. antibodies), such as in DNA-PAINT (Schnitzbauer et al. 2017), in which the specificity of a target is encoded in the DNA sequence attached to the protein label and probed by a sequence-complementary and fluorophore-labeled DNA oligonucleotide kept in the imaging buffer. This concept has the additional advantage of providing a nearly constant signal over time and being less prone to photobleaching, which has also been adapted to other super-resolution imaging techniques (Glogger et al. 2020; Spahn, Hurter, et al. 2019; Spahn, Grimm, et al. 2019).

DNA-PAINT can be extended to image multi-protein targets without requiring specialized optics in a concept termed Exchange PAINT (Jungmann et al. 2014). This method has previously been used to study multiple targets within primary neuronal cultures (Guo et al. 2019; Wang et al. 2017). However, to our knowledge DNA-PAINT has so far not been employed to study synaptic organization in neuronal tissue. Here, we have demonstrated the robustness of the Exchange PAINT method to image protein organization within the calyx of Held and principal cell in semi-thin MNTB tissue in super resolution. This method allows the imaging of multiple targets within a dense structure and is not limited by fluorophore type. Instead, the use of a single fluorophore type prevents chromatic aberration which allows the study of spatial structures with better accuracy. Furthermore, Exchange PAINT does not require the use of harsh and time-consuming elution or bleaching methods. The use of a biologically compatible imaging buffer is also advantageous in preserving target structure compared to harsh photoswitching buffers used in dSTORM imaging. Furthermore, the use of R strands speeds up image acquisition and offers exemplary image resolution and localization precision. Indeed, the resolution achieved here surpasses that achieved in similar tissue sections with dSTORM imaging by ∼5 nm (Klevanski et al. 2020). Using Exchange PAINT, multiple dense nanostructures of the pre- and post-synapse can be super-resolved to study their nanoscale spatial patterns within structurally preserved tissue sections. A possible extension would be to incorporate quantitative DNA-PAINT into this workflow, which was recently used to determine the copy numbers of AMPA receptors (Böger et al. 2019). In summary, this study reports structural cell biology with near-molecular spatial resolution using optical super-resolution microscopy.

## Methods

### MNTB tissue preparation

All experiments that involved the use of animals were performed in compliance with the relevant laws and institutional guidelines of Baden–Württemberg, Germany (protocol G-75/15). Animals were kept under environmentally controlled conditions in the absence of pathogens and *ad libitum* access to food and water. Preparation of brain sections containing the medial nucleus of the trapezoid body (MNTB) for exchange PAINT was performed according to an established protocol (Klevanski et al. 2020) with slight modifications. Briefly, Sprague-Dawley rats (Charles River) at postnatal day 13 were anaesthetized and perfused transcardially with PBS followed by 4% PFA (Sigma). Brains were dissected and further fixed in 4% PFA overnight at 4 °C. On the following day 200 µm thick vibratome (SLICER HR2, Sigmann-Elektronik) sections of the brainstem (containing MNTB) were prepared. MNTB were excised and infiltrated in 2.1 M sucrose (Sigma) in 0.1 M cacodylate buffer overnight at 4°C. Tissue was mounted on a holder, plunge-frozen in liquid nitrogen in 2.1 M sucrose and semi-thick sections (400 nm) were cut using the cryo-ultramicrotome (UC6, Leica). Sections were picked up with a custom made metal loop in a droplet of 1% methylcellulose and 1.15 M sucrose and transferred to 35 mm glass bottom dishes (MatTek) pre-coated with 30 µg/ml of fibronectin from human plasma (Sigma) and TetraSpeck fluorescent beads (1:500, Invitrogen). Dishes containing sections were stored at 4° C prior to their use.

### Antibody-DNA conjugation

Secondary antibodies of donkey anti-chicken (703-005-155), donkey anti-goat (705-005-147), donkey anti-mouse (715-005-151), and donkey anti-rabbit (711-005-152) were purchased from Jackson Immunoresearch. DNA strands were purchased from Metabion with a thiol modification on the 5’ end for each docking strand and a Cy3b dye on the 3’ end for the imager strands.

The secondary antibody to DNA docking strand conjugation was prepared using a maleimide linker as previously reported in detail (1). The thiolated DNA strands were reduced using 250 mM DTT (A39255, Thermo). The reduced DNA was purified using a Nap-5 column (17085301, GE Healthcare) to remove DTT and concentrated with a 3 kDa Amicon spin column (UFC500396, Merck Milipore).

Antibodies (>1.5 mg/mL) were reacted with the maleimide-PEG2-succinimidyl ester crosslinker in a 1:10 molar ratio and purified with 7K cutoff Zeba desalting spin columns (89882, ThermoFisher) and concentrated to >1.5 mg/mL. The DNA and antibody solutions were cross-reacted at a 10:1 molar ratio overnight and excess DNA filtered through a 100 kDa Amicon spin column (UFC510096, Merck Milipore). The antibody-DNA solution was stored at 4°C.

### Immunolabelling

Tissue samples were labelled with antibodies against α-tubulin-mouse (T6199, Sigma), TOM20-rabbit (sc-11415, SantaCruz), MAP2-chicken (188006, SySy), VGLUT1-goat (135307, SySy), Homer1/2/3-rabbit (160103, SySy), and Bassoon-mouse (SAP7F407, Enzo Life Sciences). Tissue samples in dishes were washed with PBS 3 times for 10 mins each to remove the sucrose-methylcellulose layer and blocked with 5% fetal calf serum (FCS) for 30 mins. The primary antibodies were diluted in 0.5% FCS and applied to the tissue section for 1 hour at room temperature (rt) and washed off 3x with PBS. The conjugated secondary antibody-DNA docking strand in 0.5% FCS was applied onto tissue for 1 hour at rt and washed with PBS 3x. The tissue was then stained with Alexa Fluor 488-conjugated WGA (WGA-A488) (W11261, ThermoFisher) in PBS for 10 mins and washed off 3x with PBS.

### Image acquisition

SMLM and widefield microscopy were performed on a modified Olympus IX81 inverted microscope setup with an Olympus 150x TIRF oil immersion objective (UIS2, 1.49NA) and the samples were illuminated in TIRF mode during acquisition. For imaging Cy3b DNA imager strands, a 561 nm laser line (Coherent Sapphire LP) was focused onto the sample at a density of 0.88 kW/cm^2^ through a 4L TIRF filter (TRF89902-EM, Chroma) and ET605/70 M nm bandpass filter (Chroma) and signals were detected with an Andor iXon EM+ DU-897 EMCCD camera (Oxford Instruments). WGA-A488 widefield images were obtained using a 491 nm laser line (Olympus Digital Laser System). SMLM frames were acquired using the multi-dimensional acquisition (MDA) mode in Micro-Manager 2.0 (Edelstein et al. 2014).

### Imaging conditions

DNA-PAINT imaging was performed in 5x Buffer C (2.5 M NaCl in 5x PBS) supplemented with 1mM EDTA, 1x PCA, 1x PCD, and 1x Trolox. P strands (P1 and P5) were imaged at an imager strand concentration of 0.5 nM and acquisition rate of 150 ms, and R strands (R1 and R4) at a concentration of 50 pM and acquisition rate of 100 ms. All images were acquired with 50 EM gain, for 10 000 to 20 000 frames.

### Image processing

Frames containing single molecule events were processed and rendered using Picasso software (Schnitzbauer et al. 2017). Events in each frame were localized by fitting using the Maximum Likelihood Estimation for Integrated Gaussian parameters (Smith et al. 2010). The localized events were then filtered by their width and height of the Point Spread Function (sx. sy). The resulting localizations were drift corrected using Redundant Cross-Correlation (RCC), rendered using the ‘One Pixel Blur’ function and further processed using the ‘linked localizations’ function to merge localizations with false multiple events. The NeNA value was obtained from Picasso and the decorrelation analysis to determine image resolution was calculated using an ImageJ plugin (Descloux, Grußmayer, and Radenovic 2019) based on the rendered SMLM image.

### Image analysis

The length of Bassoon and Homer were measured in ImageJ by creating a binary mask of the rendered image with the preset ‘moments’ threshold. A line was drawn along the long axis of the AZ and PSD structure, respectively, and the length was measured. The distance between Bassoon and Homer was measured by drawing a line perpendicular to both structures and adjusting the spline fit to incorporate the linear length of the structures. The fluorescence intensity for each structure was plotted and fitted with a Gaussian function. The distance was calculated from the distance between the peak intensities of the two structures. Similarly, the line profile of Bassoon, Homer, and VGLUT was obtained by measuring their fluorescence intensity using the line tool with spline fit perpendicular to the structures. Fluorescence intensity against distance was averaged for all ROIs with Bassoon peak intensity as the reference point.

FRC analysis was performed by saving filtered and drift-corrected DNA-PAINT localizations from Picasso and opening the localizations in ThunderSTORM (Ovesný et al. 2014). Localizations were filtered according to frame length from 0 to n and each frame length was filtered into blocks of 100. Rendered images were saved and FRC values were calculated using the BIOP.FRC plugin in ImageJ with the Fixed 1/7 criteria.

### Sequences of DNA oligonucleotides

**Table 1:** Sequences of docking strands

| Name | Sequence | Modification |
| --- | --- | --- |
| P1 docking strand | TTATACATCTA | 5' - Thiol |
| P5 docking strand | TTTCAATGTAT | 5' - Thiol |
| R1 docking strand | TCCTCCTCCTCCTCCTCCT | 5' - Azide |
| R4 docking strand | ACACACACACACACACACA | 5' - Azide |

**Table 2:** Sequences of imager strands

| Name | Sequence | Modification |
| --- | --- | --- |
| P1 imager strand | TAGATGTAT | 5' - Cy3B |
| P5 imager strand | CATACATTGA | 5' - Cy3B |
| R1 imager strand | AGGAGGA | 5' - Cy3B |
| R4 imager strand | TGTGTGT | 5' - Cy3B |

## Acknowledgements

M.H. and T.K. acknowledge funding by the Baden-Württemberg Foundation (Mult!Nano, Methods in life sciences program), in whose name this research was conducted. M.H., N.D-H., M.D. and Y.L. acknowledge funding by the Deutsche Forschungsgemeinschaft (DFG, grant SFB 902) and the Volkswagen foundation (grant 91067-9). We are grateful to Christoph Spahn for support with FRC analysis.

## Notes

### Competing Interest Statement

The authors have declared no competing interest.

